# Antidiabetic Drugs May Prevent Kidney Damage

**DOI:** 10.1101/2023.10.19.563071

**Authors:** Maria de Fatima Fernandes Vattimo, Eloiza Oliveira Silva, Sara Ventura, Camila Lima, Jessica Garcia, Alessandra Oliveira Maia, Juliana Veloso Gusmão, Guilherme Henrique Ferreira Vieira, Carla Djamila de Pina Victoria

## Abstract

**Introduction:** Sodium-glucose cotransporter 2 inhibitors (SGLT2i) are a class of drugs that act as glucose reducers in patients with type 2 diabetes mellitus. Recent meta-analyses have shown that SGLT2i also prevent acute kidney injury (AKI) in diabetic patients. The aim of this study was to investigate the protective effect of canagliflozin an SGLT2i on AKI due to ischemia and reperfusion (I/R) in non-diabetic rats.

**Methods:** Male Wistar rats weighing 250-300 g were divided into four groups: Control; SHAM (rats submitted to surgical simulation of renal ischemia); I/R: rats submitted to renal ischemia (bilateral clamping of the renal hilum for 30 minutes); CANA+I/R: I/R rats that received canagliflozin (30mg/kg, oral, gavage; 5 days before I/R). Renal function parameters were evaluated (serum creatinine [CrS], inulin clearance [Clin]; renal hemodynamics mean arterial pressure [MAP], renal blood flow [RBF], renal vascular resistance [RVR]); oxidative profile (urinary peroxides - FOX, lipid peroxidation - TBARS, urinary nitrate- NO) and thiols in renal tissue and expression of nuclear factor-erythroid 2 related factor 2 [Nrf2] protein.

**Results:** The I/R group showed an increase in CrS and a reduction in inulin clearance, while the CANA+I/R group showed a reduction in serum creatinine and an increase in inulin clearance compared to the I/R group. In addition, the CANA+I/R group showed a decrease in oxidative metabolites (FOX and TBARS) and an increase in Nrf2 compared to the I/R group.

**Conclusion:** Canagliflozin treatment prevented the reduction in renal function induced by ischemia and reperfusion. In addition, there was a reduction in oxidative activity due to a decrease in oxidative metabolites and urinary peroxides and an increase in renal tissue thiols and Nrf-2, which is responsible for transcribing antioxidant activity. Therefore, the current study confirmed a relevant renoprotective effect of canagliflozin in the presence of renal ischemia.

## INTRODUCTION

Acute kidney injury (AKI) is a syndrome characterized by a sudden decline in kidney function. ^(1)^ Its classifications include pre-renal and renal, which can be associated with ischemic events such as sepsis, shock, infections, the use of radiological contrasts, and drug toxicity.^(2)^

According to Kidney Disease: Improving Global Outcomes (KDIGO), 1 in 5 adults (21.6%) and 1 in 3 children (33.7%) have AKI worldwide. Clinically, this syndrome is defined as an increase in serum creatinine of 0.3mg/dL in 48 hours, an increase of 1.5 times the baseline value in 7 days, or a reduction in urinary flow to less than 0.5mL/Kg for 6 hours.^(3)^

It is known that the kidneys receive around 20% of the body cardiac output, making them highly vulnerable to reductions in perfusion and oxygenation. In a severe or prolonged reduction in blood flow or renal hypoxia, cellular integrity and metabolism are compromised, ultimately leading to vascular, glomerular, and tubular dysfunction. The damage is characterized by the glomerular filtration rate (GFR), which decreases in the event of injury. However, GFR does not always correlate with the patient’s underlying condition, demonstrating the complexity of vascular and tubular processes in renal dysfunction.^(4)^

The pathophysiology of AKI can be described as a series of events that range from acute injury to gradual onset over long periods of time, as in episodes of chronic kidney disease. These events trigger compensatory mechanisms in the kidneys, such as initial hyperfiltration and hypertrophy of the remaining viable nephrons. As this involvement persists, subsequent histopathological changes follow, including distortion of the glomerular architecture, abnormal podocyte function, and interruption of filtration leading to renal sclerosis.^(5)^

This syndrome affects 30%-60% of critically ill patients and is associated with acute morbidity and mortality.^(6)^ In addition, there is evidence showing that a large proportion of patients affected by AKI progress from the acute phase to chronic kidney disease (CKD), and accelerated progression to end-stage renal disease (ESRD), increased risk of cardiovascular complications, recurrent episodes of AKI, and mortality^(7)^ leading to poor quality of life, disability, and high long-term therapeutic costs.^(8)^

Studies seek to mitigate the complications of AKI in patients whose hemodynamic and immunological reserves are restricted. ^(9,10,11)^ Treatments that are still under discussion are needed to reduce high morbidity and mortality and improve the recovery of kidney function. In some cases, such as preoperative risk for kidney disease or surgical risk, there is an opportunity to implement preventive measures to preserve kidney function. Alternatively, it could de possible to intervene after the patient has been diagnosed with AKI to try to improve recovery and reduce progression to CKD / ESRD in high-risk patients.^(12)^

One of the most widely used pharmacological therapies for controlling the progression of kidney disease is angiotensin-converting enzyme inhibitors. However, recent clinical trials have evaluated the impact of sodium-glucose cotransporter 2 inhibitors (SGLT2i), which demonstrate cardiovascular and renal benefits, suggesting that they are independent of the impact on glycemia. ^(13)^

Gliflozins, SGLT2i, cause a reduction in glycated hemoglobin without causing hypoglycemic effects, as well as natriuresis and consequent volume reduction. The SGLT2 co-transporter is located in the proximal convoluted tubule of the S1 nephron and is responsible for reabsorbing 90% of the filtered glucose load. Therefore, SGLT2i blocks the renal reabsorption of glucose and sodium by reducing the absorption of sodium and glucose in the proximal convoluted tubule. A tubuloglomerular feedback occurs, which increases the arrival of sodium in the distal convoluted tubule, causing an impact on the macula densa and consequent vasoconstriction of the afferent arteriole, leading to glycosuria and natriuresis, which contributes to a reduction in intraglomerular pressure (glomerular hyperfiltration) and hemodynamic stress, suggesting one of the SGLT2i mechanisms of renoprotection.^(13)^ In addition, they are associated with a substantial reduction in albuminuria, pro-inflammatory mediators, and predominant changes in renal hemodynamics.^(14)^

Studies with animal models of ischemia/reperfusion allow the transposition of clinical situations to the bench, in an effort to isolate variables that cannot be approached in humans, and identify pathophysiological mechanisms and physiological responses to pharmacological or non-pharmacological therapies and that are still inconclusive. This reinforces the need for studies on the effects and renal mechanisms of anti-diabetic drugs, glyphozines. Therefore, the hypothesis of this study is that these agents play a protective role in renal function and renal oxidation in AKI.

The aim of this study was to evaluate the effect of the antidiabetic drug canagliflozin on the function, hemodynamics, oxidative profile, and histology of rats submitted to a model of acute ischemic renal injury.

## MATERIALS AND METHODS

### Ethical aspects

This quantitative animal model study was approved by the Ethics Committee for the Use of Animals at the University of São Paulo School of Medicine (CEUA - FMUSP) under registration no. 1624/2021. All procedures carried out in this study were in accordance with the Ethical Principles for Animal Experimentation adopted by the Brazilian College of Animal Experimentation.^(15)^

The sample size was determined using G*Power software, using the one-way ANOVA variance test (F-test). The subjects were 20 adult male Wistar rats, 60 days old, weighing 250 to 300 g. They were kept with free access to water and food, in suitable thermal conditions, and in alternating day and night cycles. They were randomly allocated into 4 (four) groups: SHAM (n = 5) - animals submitted to laparotomy to simulate clamping of the bilateral renal pedicles for 30 min; CANA group (n=10) - animals that received canagliflozin by gavage (200 mg/kg, once a day, for 5 days, simulating renal ischemia); Ischemia (I/R) group (n = 5) - animals submitted to laparotomy to clamp the bilateral renal pedicles for 30 min, followed by reperfusion; CANA + Ischemia group (n = 10) - animals given canagliflozin by gavage (200 mg/kg, 1× a day, orally, for 4 days. On the 5th day, they underwent renal ischemia)

After 24 hours of post-anesthetic recovery, the animals were placed in metabolic cages to collect 24-hour urine to study renal function and oxidative stress. All animals were evaluated during anesthetic recovery and received post-procedure analgesics (tramadol 15 mg/kg intramuscular 3x a day for 2 days). After this period, the rats were removed from their cages, anaesthetized, and subjected to renal function analysis using the inulin clearance technique.

The animals underwent laparotomy and terminal blood collection by puncture of the abdominal aorta. The left kidney was removed and stored in a refrigerator at -80 °C to check for nonprotein thiols. At the end of the experiment, the researcher euthanized the animals by collecting terminal blood, in accordance with the ethical standards for handling animals in laboratories.

The glomerular filtration rate (GFR) was determined using the inulin clearance technique. Preoperatively, the animals received analgesia with morphine (3 mg/kg) and were anesthetized with ketamine and xylazine (100 mg/kg and 10 mg/kg, respectively). Once anesthetized, the operation began with catheterization of the animal’s jugular vein for bolus infusion of the initial dose of 100 mg/kg of body weight of diluted inulin, followed by continuous infusion of 10 mg/kg of body weight during the 2 hours of the experiment, at a rate of 0.04 mL/min. Urine was collected every 30 minutes by bladder catheterization and blood samples every 60 minutes. The animals’ sedation was maintained with an additional dose of xylazine, if necessary. ^(16)^

Urinary and plasma creatinine levels were determined using the colorimetry method known as the Jaffé method. ^(17)^

Renal hemodynamics was checked using renal blood flow (RBF) with an ultrasonic probe (T402; Transonic Systems, Bethesda, MD, USA), which involved the left renal artery. Renal vascular resistance (RVR) was calculated using the following formula: RVR = mean arterial pressure (MAP)/RBF. Measurement of MAP was carried out by catheterization of the carotid artery for insertion of the catheter (polyethylene tube - PE 60), and terminal blood was collected by puncture of the abdominal aorta, and the kidneys were sectioned for histopathological study. ^(18)^

The oxidative profile was assessed by measuring urinary peroxides (UP) using the FOX-2 method.^(19)^ The end products of lipid peroxidation were detected by the urinary thiobarbituric acid reactive substances method (TBARS).^(20)^ Urinary nitrate was quantified by the Griess method^(21)^ and non-protein soluble thiols in renal tissue with correction for total proteins were quantified by the Ellman method.^(22)^ Proteins were extracted from mouse renal tissues and were subjected to Western blot analysis after quantification using the Bradford method. After incubating with primary antibodies at 4°C overnight, the samples were exposed to secondary antibodies for 60 min at room temperature. Protein expressions were detected using the Odyssey infrared imaging system from LICOR (U.S.A.), and the data were analyzed and normalized to the corresponding β-actin. Primary antibody of NRF2 (MA542371) were purchased from Termo Fisher Scientific, Brasil. In renal histology, to evaluate tubulointerstitial involvement, a score graded on a scale of 0 to 4 was used [20], where 0 = normal; 0.5 = small focal areas; 1 = involvement of less than 10% of the cortex and external renal medulla; 2 = involvement of 10% to 25% of the external renal cortex; 3 = involvement of 25% to 75% of the cortex and external renal medulla; and 4 = extensive alterations of more than 75% of the cortex and external medulla. The images obtained by optical microscopy were captured using a lightweight video camera connected to an image analyzer and examined using fields of 0.245 mm, for 2 slides containing a sample of kidney tissue from each animal.

The data was analyzed using one-way ANOVA. A significance level of p < 0.05 was chosen, considering that the effect of at least one of the groups was different from the others. Tukey’s 2 by 2 multiple comparisons tests were used to evaluate which groups differed from each other.

## RESULTS

In the results presented below, the animals in the SHAM group were used as healthy controls and the data obtained for this group were considered normal parameters.

### Renal function

Renal function was evaluated using the parameters of urinary flow, serum creatinine, and inulin clearance, with the SHAM group presenting values that were considered to be normal, as shown in Table 1.

**Table 1.**
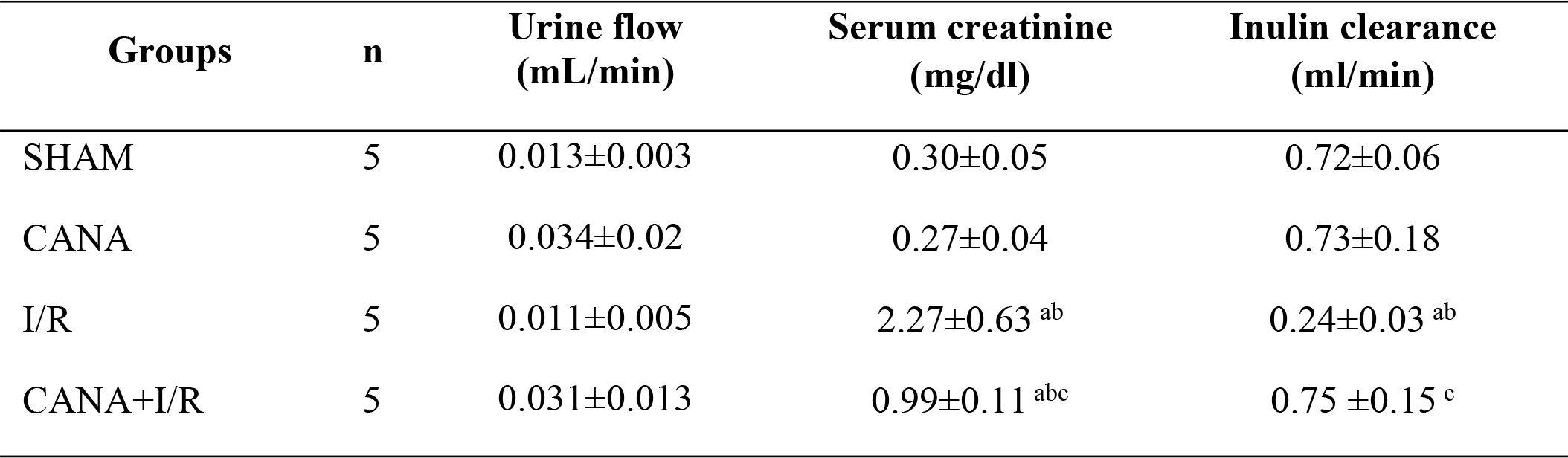
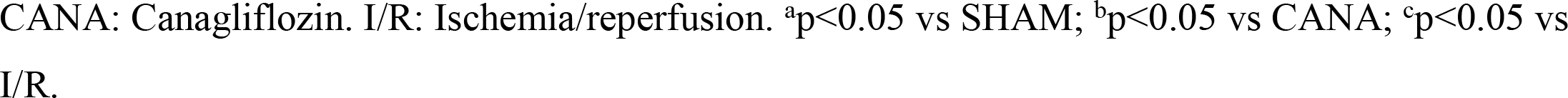
Renal function.

| Groups | n | Urine flow<br>(mL/min) | Serum creatinine<br>(mg/dl) | Inulin clearance<br>(ml/min) |
| --- | --- | --- | --- | --- |
| SHAM | 5 | 0.013±0.003 | 0.30±0.05 | 0.72±0.06 |
| CANA | 5 | 0.034±0.02 | 0.27±0.04 | 0.73±0.18 |
| I/R | 5 | 0.011±0.005 | 2.27±0.63<sup>ab</sup> | 0.24±0.03<sup>ab</sup> |
| CANA+I/R | 5 | 0.031±0.013 | 0.99±0.11<sup>abc</sup> | 0.75 ±0.15<sup>c</sup> |

CANA: Canagliflozin. I/R: Ischemia/reperfusion. <sup>a</sup>p<0.05 vs SHAM; <sup>b</sup>p<0.05 vs CANA; <sup>c</sup>p<0.05 vs I/R.

In the CANA group, there was an increase in urinary flow compared to the SHAM group. In terms of serum creatinine and inulin clearance, there was no statistical difference between the CANA and SHAM groups.

The I/R group showed a decrease in urinary flow and inulin clearance and an increase in serum creatinine when compared to the SHAM and CANA groups.

The CANA + I/R group showed an increase in urinary flow when compared to the SHAM and I/R groups. CANA+I/R group showed an increase in serum creatinine when compared to the SHAM and CANA groups, however, when compared to the I/R group, it showed a reduction in this parameter. SHAM and CANA groups showed no difference in inulin clearance when compared to each other and CANA+I/R presented an increase in inulin clearance, similar to the values showed by the control groups, when compared to the I/R group, confirming the renoprotective effect of the canagliflozin in this group.

### Renal hemodynamics

Renal hemodynamics were evaluated by checking renal blood flow (RBF) and renal vascular resistance (RVR) as shown in Table 2. The renal blood flow of the SHAM and CANA groups were statistically similar.

**Table 2.** Renal hemodynamics.

| Groups | n | Renal blood flow<br>(mL/min) | Renal vascular<br>resistance<br>(mmHg/ml/min) |
| --- | --- | --- | --- |
| SHAM | 5 | 9.28±1.73 | 11.10±1.6 |
| CANA | 5 | 7.42±1.52 | 12.19±2.98 |
| I/R | 5 | 1.61±0.26<sup>ab</sup> | 56.77±0.26<sup>ab</sup> |
| CANA+I/R | 5 | 6.80±0.60<sup>ac</sup> | 13.27±1.61<sup>c</sup> |

CANA: Canagliflozin. I/R: Ischemia/reperfusion. <sup>a</sup>p<0.05 vs SHAM; <sup>b</sup>p<0.05 vs CANA; <sup>c</sup>p<0.05 vs I/R

I/R group showed a significant reduction in RBF when compared to the SHAM and CANA control groups, while the CANA+I/R group showed a significant increase in this parameter compared to the I/R group. SHAM and CANA control groups showed normal values with no statistical difference in the RVR. The I/R group showed an additional significant increase in RVR compared to the SHAM and CANA groups, while the CANA+I/R group showed a significant reduction in this parameter compared to the I/R group.

### Oxidative profile

As shown in Table 3, the control groups (SHAM and CANA) were considered normal and showed no significant differences between them for any of the parameters analyzed.

**Table 3.** Oxidative profile of animals.

| Groups | n | Urinary peroxides<br>(nmol/g urine<br>creatinine) | Lipid peroxidation<br>(mnol/g urine<br>creatinine) | Urinary nitrate<br>(nmol/g urine<br>creatinine) | Thiols (nmol/mg total<br>protein) |
| --- | --- | --- | --- | --- | --- |
| SHAM | 5 | 3.30±0.6 | 0.18±0.01 | 16.00±5.20 | 15.70±6.1 |
| CANA | 5 | 4.18±1.41 | 0.10±0.21 | 36.75±6.41<sup>a</sup> | 16.02±7.31 |
| I/R | 5 | 11.64±4.00<sup>ab</sup> | 0.43±0.07<sup>ab</sup> | 174.51±13.02<sup>ab</sup> | 1.70±0.24<sup>ab</sup> |
| CANA+I/R | 5 | 6.68±1.18<sup>c</sup> | 0.50±0.07<sup>ac</sup> | 144.56 ±48.78<sup>ab</sup> | 12.61±19.39<sup>abc</sup> |

CANA: Canagliflozin. I/R: Ischemia/reperfusion. <sup>a</sup>p<0.05 vs SHAM; <sup>b</sup>p<0.05 vs CANA; <sup>c</sup>p<0.05 vs I/R

**Figure 1.**
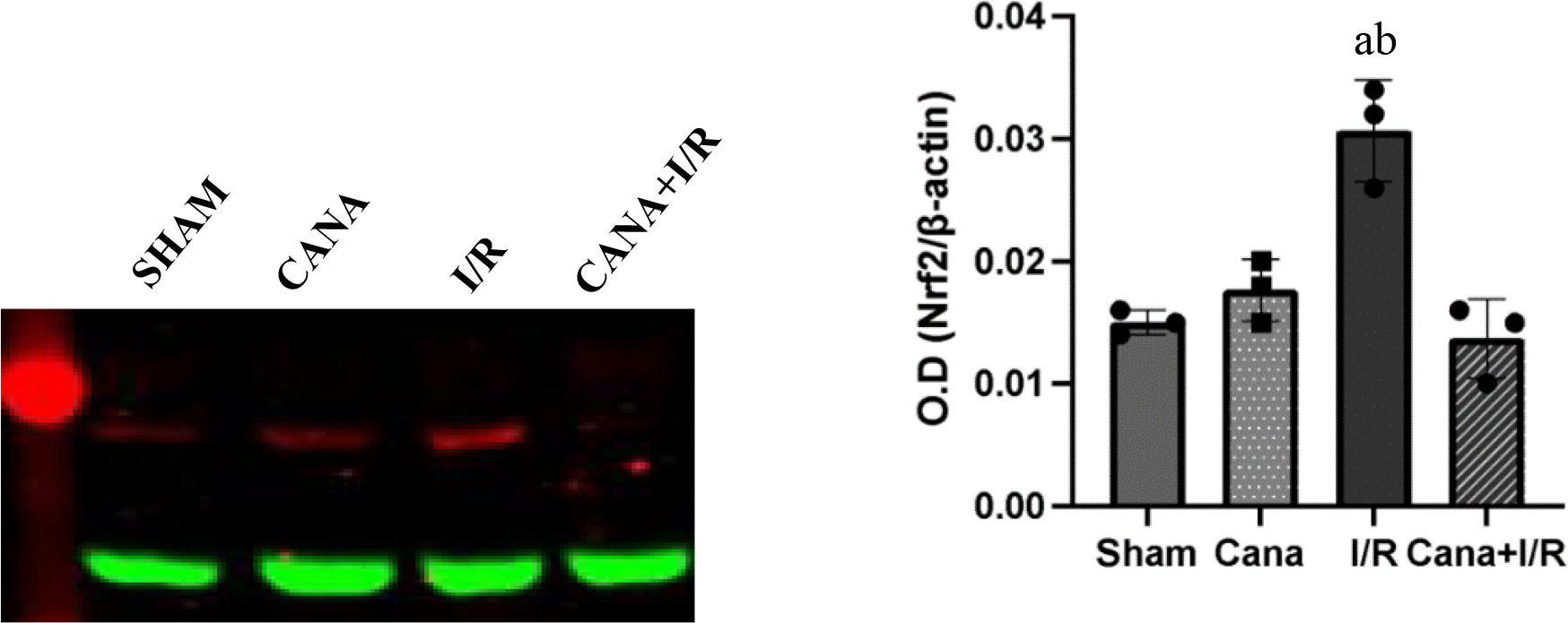
Erythroid 2 factor NRF2. CANA: Canagliflozin. I/R: Ischemia/reperfusion. ^a^p<0.05 vs SHAM; ^b^p<0.05 vs CANA; ^c^p<0.05 vs I/R

The CANA+I/R group showed a reduction in the levels of urinary peroxides, lipid peroxidation and urinary nitrate when compared to the I/R group. The I/R in the untreated group led to an additional increase in these oxidative metabolites when compared to the SHAM and CANA control groups.

I/R group showed a significant reduction in the levels of thiols in the kidney tissue when compared to the control groups. The CANA+I/R group showed a significant increase in this parameter when compared to the I/R group.

The SHAM and CANA groups showed a significant reduction in this parameter. However, NrF2 transcription was elevated in the I/R group when compared to the other groups. The CANA+I/R group showed a significant reduction in this parameter when compared to the I/R group.

## DISCUSSION

The present study evaluated the effects of canagliflozin, or glyphlozin, in rats submitted to the I/R-induced renal failure model. The results confirmed the renoprotective effect of this antidiabetic agent in the renal function of animals submitted to I/R AKI.

The complications of renal function resulting from I/R were demonstrated in the experimental model, which consisted of transient hypoperfusion lasting 30 minutes due to clamping of the renal pedicles, triggering a response to renal ischemia. This finding was confirmed by the results of renal function parameters and oxidative stress already described to the injury.

I/R injury involves several pathophysiological mechanisms, damage and dysfunction in the endothelial and tubular cells, an inflammatory response, and the formation of reactive oxygen species.^(23)^ These animals showed impaired renal function, which was demonstrated by reduced creatinine clearance, elevated serum creatinine, and intensification of the redox mechanism, corroborating the findings of other experimental studies using the same acute injury model.^(24)^

The tubular lesions aggravated during I/R are most likely related to mechanisms of reactive oxygen species (ROS) production, once the animals subjected to I/R had showed an increase in oxidative metabolites (urinary peroxides and TBARS). Hydrogen peroxides, whose urinary excretion was shown to be high in I/R animals, contribute to the occurrence of lipid peroxidation and consequent structural and functional damage to the affected cells.^(25)^

Canagliflozin conditioning was associated with an improvement in renal function parameters, which was confirmed by an increase in creatinine clearance, a decrease in serum creatinine, and a reduction in urinary flow. In this study, the measurement of serum creatinine and creatinine clearance in the group only receiving the drug, CANA group, was similar to the control group. In addition, there was modulation of renal hemodynamics, corroborating other studies. This indicates that canagliflozin may have an impact on improving renal function by improving renal hemodynamics, and can thus could be used to prevent ischemic acute kidney injury.^(26,27)^

Given the worsening of oxidative stress in I/R, canagliflozin triggered an improvement in oxidation-related parameters, evidenced by a reduction in oxidative metabolites and an increase in the thiol pool. Interestingly, Nrf2 activity was reduced in the animals in the CANA+I/R group, suggesting that canagliflozin has a suppressive effect on Nrf2 activity during I/R. This reduced of Nrf2 is related to the absence of oxidative stress conditions in these animals.^(28,29)^ The NRF2 is associated with a molecule called Keap 1, when Keap1 is oxidized this molecule releases NRF2 leaving it free to be translocated to the cell nucleus and act as a transcription factor, activating antioxidant response^(29)^.

In addition to its efficacy as a glucose reducer in patients with type 2 diabetes mellitus, Sodium-glucose cotransporter 2 inhibitors, such as canagliflozin, have demonstrated an intriguing antioxidative mechanism. Recent studies indicate that canagliflozin exerts a protective effect against AKI in diabetic patients. This additional benefit can be attributed to canagliflozin’s ability to modulate oxidative processes, thereby reducing oxidative stress and preserving renal function. The antioxidant property of canagliflozin represents a significant advancement in the therapeutic arsenal available for the treatment of patients with AKI and holds promising prospects for the care of their kidneys^(30)^.

Studies corroborate that SGLT2i promotes anti-inflammatory, antioxidant, and antifibrotic actions, improves renal oxygenation, and reduces glomerular hypertension and hyperfiltration. Experimental models have linked SGLT2i to reductions in oxidative stress using markers of inflammation (nuclear factor kB, interleukin 6, monocyte, chemoattractant protein-1, macrophage infiltration) and fibrosis (fibronectin, transforming growth factor-b) and attenuated histological evidence of nephropathy at the structural level. SGLT2 inhibition reduces glomerular and interstitial tubule damage in several animal models of diabetes.^(30,31)^

SGLT2 inhibitors also reduce renal ischemic injury. Proximal tubular oxygen supply is related to changes in hematocrit. Data suggests that the increase in hematocrit with SGLT2 inhibition is due to the normalization of renal cortical oxygenation, restoring normal cellular function to the erythropoietin-producing cells in the kidney, which increases hematocrit levels.^(32)^

Therefore, this study proved the efficacy of SGLT2i as a renoprotective agent in the presence of I/R. These findings suggest that the protective role of pretreatment with canagliflozin was achieved mainly by modulating renal hemodynamics, showing it to be a promising treatment for patients at risk of developing AKI.

## CONCLUSION

Therapy of male Wistar rats that were submitted to the renal ischemia model with canagliflozin resulted in the preservation of renal function, since preconditioning with the drug prevented AKI, as evidenced by the equivalence of inulin clearance between the control and treated groups, and also by the increase in serum creatinine in the untreated group compared to the group that underwent I/R and received the drug.

The antioxidant activity of the drug is also shown by the decrease in oxidative metabolites and increase in antioxidants, which may have contributed to a reduction in glomerular and tubular damage. The results corroborate the therapeutic benefits of this drug in preventing AKI in the animal model.

### Research funding

This research was funded by the São Paulo Research Foundation (FAPESP), No. 2019/03415-3, and the Coordination for the Improvement of Higher Education Personnel - Brazil (CAPES), Financial Code 001.

